# Correction of copy number induced false positives in CRISPR screens

**DOI:** 10.1101/151985

**Authors:** Antoine de Weck, Javad Golji, Mike Jones, Joshua Korn, Eric Billy, E. Robert McDonald, Tobias Schmelzle, Hans Bitter, Audrey Kauffmann

**Affiliations:** Novartis Institutes for Biomedical Research, Basel CH-4002, Switzerland.; Novartis Institutes for Biomedical Research, Cambridge, MA 02139, USA.

## Abstract

Cell autonomous cancer dependencies are now routinely identified using CRISPR loss-of-function screens. However, a bias exists that makes it difficult to assess the true essentiality of genes located in amplicons, since the entire amplified region can exhibit lethal scores. These false-positive hits can either be discarded from further analysis, which in cancer models can represent a significant number of hits, or methods can be developed to rescue the true-positives within amplified regions. We propose two methods to rescue true positive hits in amplified regions by correcting for this copy number artefact. The Local Drop Out (LDO) method uses the relative lethality scores within genomic regions to assess true essentiality and does not require additional orthogonal data (e.g. copy number value). LDO is meant to be used in screens covering a dense region of the genome (e.g. a whole chromosome or the whole genome). The General Additive Model (GAM) method models the screening data as a function of the known copy number values and removes the systematic effect from the measured lethality. GAM does not require the same density as LDO, but does require prior knowledge of the copy number values. Both methods have been developed with single sample experiments in mind so that the correction can be applied even in smaller screens. Here we demonstrate the efficacy of both methods at removing the copy number effect and rescuing hits from some of the amplified regions. We estimate a 70-80% decrease of false positive hits in regions of high copy number with either method.

## Introduction

CRISPR based loss-of-function screens have emerged as a powerful tool to interrogate multiple species and models (1). The technology has been quickly adopted to identify essential genes in cancer, including several cancer cell line screens (2–4). However, as reported in two studies (5,6) and further discussed by others (7), genes in regions of copy number amplification display strong lethal phenotypes by CRISPR-Cas9 cutting (as opposed to CRISPRi (8)), regardless of the true biological essentiality of the targeted gene. This results in a significant number of false positive hits in samples with large copy number alterations. This is of particular relevance in cancer models, which typically display extensive copy number events.

One way of mitigating the problem of false positives would be to simply discard any hits found in amplified regions. This is a viable strategy when considering aggregate profiles (9), but runs the risk of yielding many false negatives when looking at individual hits. Especially when copy number events are an important oncogenic driver and identifying the essential gene in the amplicon is of interest to target discovery (10). Therefore, to fully leverage CRISPR based screens, it is important to understand and correct for the observed copy number bias. Here, we propose methods to correct for the copy number artefact, while rescuing the true positives within the amplicons. The corresponding R scripts are also provided (https://doi.org/10.6084/m9.figshare.5140057.v1).

We used the data published by Munoz et al. (5), where the copy number artefact has been observed (Fig. 1), i.e. a negative correlation of sensitivity (calculated as Log FC) with copy number. To illustrate the methods, we focused on the astrocytoma cell line sf268 and the gastric cancer cell line mkn45, as these two cell lines have amplicons where the driver has been well characterized, *YAP1* and *MET*, respectively. The sgRNA library used targeted 2722 human genes with an average coverage of 20 reagents per gene. In addition, a second screen performed on mkn45, using a different library of genes with a coverage of 10 reagents per gene, was used to evaluate the methods described herein.

**Figure 1.**
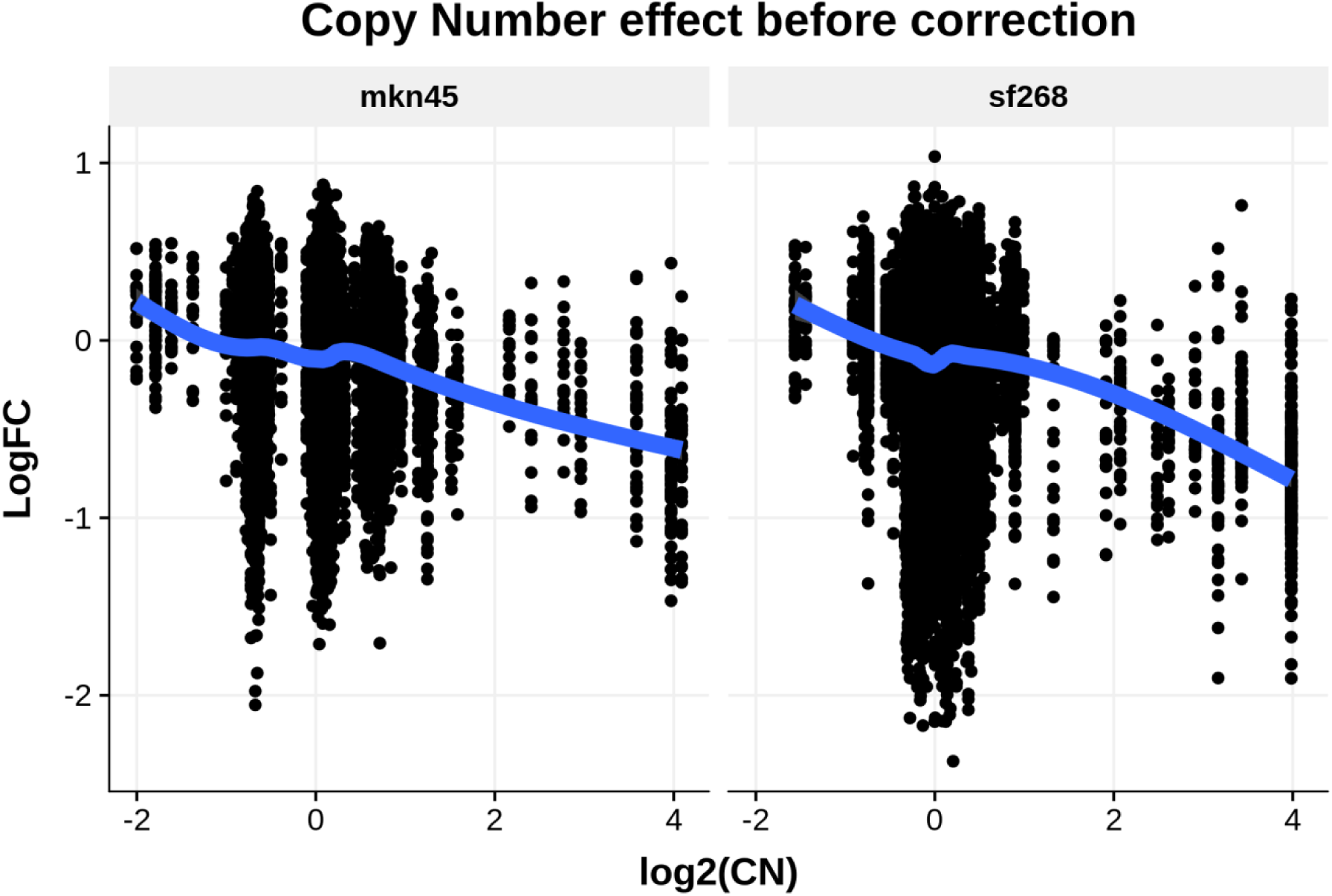
CN effect on CRISPR knock-out sensitivity. The sensitivity to CRISPR-mediated knock-out is dependent on the level of amplification of the underlying genomic region. Above for mkn45, 84 out of 191 guides in amplified regions (CN of at least 4 (log2(CN)=2)) score below -0.5, while 274 out of 397 guides score below -0.5 in sf268.

### Local Drop Out (LDO) method

To account for the copy number artefact, we propose the Local Drop Out (LDO) method. LDO aims to correct phenotype scores for each guide by taking into account guide scores targeting the other genes in its direct genomic neighbourhood. It assumes that most genes display little or no phenotype upon knock-out in such screens (˜2 weeks or less) and does not rely on copy number measurements. If multiple neighbouring genes show similarly strong drop out values, it is assumed that the observed phenotype is due to a copy number effect rather than a true dependence of the cell line. The density of the screen influences the size of the copy number events that can be detected: the higher the density of the genes selected to be included in the screen, the more focal the detected copy number events can be.

The LDO method uses a two-step process. First, LDO aims to identify as many guides as possible that may be true positives, which include both essential genes and growth enhancers. This is done in order to remove them from consideration for the second step and maximize the proportion of “neutral” genes used to estimate the CN specific effect. Although not required, prior knowledge can be used, e.g. lists of pan-lethal genes available in the public domain can be leveraged to determine an initial list of essential guides for consideration. Additional cell line specific essential or growth enhancing genes can be identified by calculating, for each guide, a weighted mean sensitivity of neighbouring guides. The weighting function is parameterized in the calculation. To identify additional essential or growth enhancing genes, we used an exponential distribution with parameter *ω* = 100’000 bp and performed the calculations independently for each chromosome. The guides targeting the same gene were not included in the calculation; neither were the guides targeting genes known to be essential as per the pan-lethal list. The latter group of guides was removed since we aimed to estimate the CN effect on only the guides whose drop out can only be attributed to the artefactual CN effect. Any guide targeting an essential gene is expected to display a true phenotype in addition to any potential CN effect and their inclusion is therefore likely to partially hide the signal we aim to detect. In this analysis, we have used a list of a priori essential genes compiled from [4] (see Mat & Met for more details). Removing the known pan-essential genes is however not a requirement of the method, but can improve the accuracy of the resulting CN correction. In particular in the case of successively located pan-essential genes which could otherwise be confused for a CN event.

The weighted mean sensitivity is calculated as follows. Let *g* be a guide in the set *G* of all guides under consideration (e.g. all guides in a specific chromosome or chromosomal arm), with 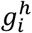 the *i*th guide targeting gene *h* and *G^h^* be the set of guides targeting gene *h*. Let *E*_1_ be the set of guides targeting known essential genes.Additionally, let the genomic position of guide *g* be *x_g_* and the sensitivity induced by guide *g* be *S_g_*, then the weighted mean sensitivity, excluding essential guides and guides targeting the same gene, 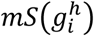 for guide 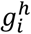 can be written as:

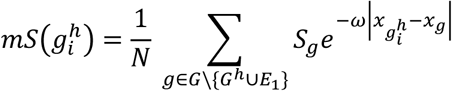

With

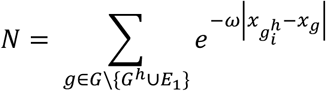

For each guide, the first iteration of the corrected sensitivity *S*^1^ value is obtained from subtracting the weighted mean sensitivity for that guide to the original sensitivity value without correction (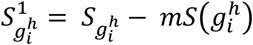). Using this measure, the guides with absolute values above the *μ^th^* percentile across the entire genome are considered as guides displaying potential true phenotypes and are not used in the second iteration of the method (by default *μ* = 85). Here *μ* = 85 is chosen to represent a prior belief that we can expect about 15% of genes (and therefore 15% of guides in our design) will display a true phenotype in the screen independent of the copy number. The parameter can be modified, e.g. one might expect a larger percentage of genes displaying a phenotype in longer screens. This procedure is equivalent to increasing the set of essential guides in set *E*_2_ which is then sample specific and contains the set *E*_1_ and all the guides identified above the *μ^th^* percentile.

In the second step of the LDO correction, all guides below the *μ^th^* percentile are used to fit two regression trees to estimate the copy number effect. The first regression tree is parametrized to capture short amplifications while the second regression tree is optimized to capture large chromosomal arm events. These guides are highly enriched in guides showing no phenotypes, i.e. the set of guides *g* ∊ *G*\*E*_2_. Here, one dimensional regression trees are used to estimate the sensitivity of the guides as a function of the genomic location alone. The combined copy number effect of the two regressions is then removed from the original sensitivity score *S_g_* to obtain the LDO corrected sensitivity score 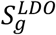. Specifically, the regression tree *T* formulates the copy number induced sensitivity *S_CN_* at position *x* as follows:

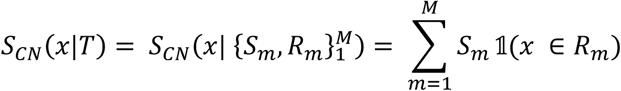

Where 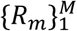 are subregions of the genome, and *x* is a genomic position. *S_m_* are the estimated copy number induced sensitivity values in region *R_m_*. Using only the guides ∊ *G*\*E*_2_, we try to find the regression tree *T* which minimizes the error:

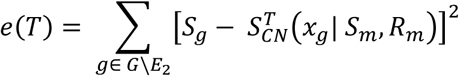

with respect to *S_m_* and *R_m_*. In practice, a regularization term is added to avoid overfitting. Thus, the objective is to identify the tree *T* which minimizes the following term:

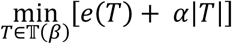

where |*T*| is the number of terminal nodes of the tree and the complexity parameter *α* measures the “cost” of adding another region *R_m_* to the model. The higher the cost, the shallower the tree. Also additional constraints can be set on the universe 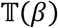 of potential trees *T*. In particular, one can consider the universe 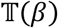 with a minimum number *β* of guides per region *R_m_*.

The first regression performed attempts to identify short focal amplifications. As a result, relatively loose constraints are applied: the default parameters chosen were *α*_1_ = 0.001 and *β*_1_ = 1.5 times the mean number of guides per gene. Furthermore, only the regions 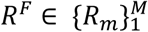 with at least 3 genes and with correction values |*S_m_* | > 1.5 *mad* (*S_g_* — *S_CN_*(*x_g_*)) with *mad* = median absolute deviation are considered as true regions of focal copy number events.

The second regression sets out to identify large chromosomal abnormalities from the remaining regions 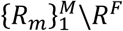. Here, tighter constraints are used, specifically *α*_2_ = 0.01 and *β*_2_ = 10% of the number of guides in *G*. The resulting regions with their respective correction scores are combined with the regions identified in the focal amplification regression tree to build the final LDO tree *T^LDO^*, i.e. the union of the disjoint first (*R^F^*) and second tree 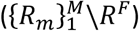 is considered. It follows that the LDO corrected sensitivity scores is defined by:

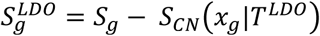

In Fig. 2, the essentiality score before and after LDO correction is shown for the *YAP1* amplicon in sf268. In Fig. 2 a), *ANGPTL5*, *KIAA1377*, *C11orf70*, *BIRC3*, *BIRC2*, *TMEM123*, *MMP7*, and *MMP20* display equivalently significant phenotypes believed to be entirely due to the copy number artefact. On the other hand, YAP1 shows a stronger phenotype relative to its neighbouring genes.

**Figure 2.**
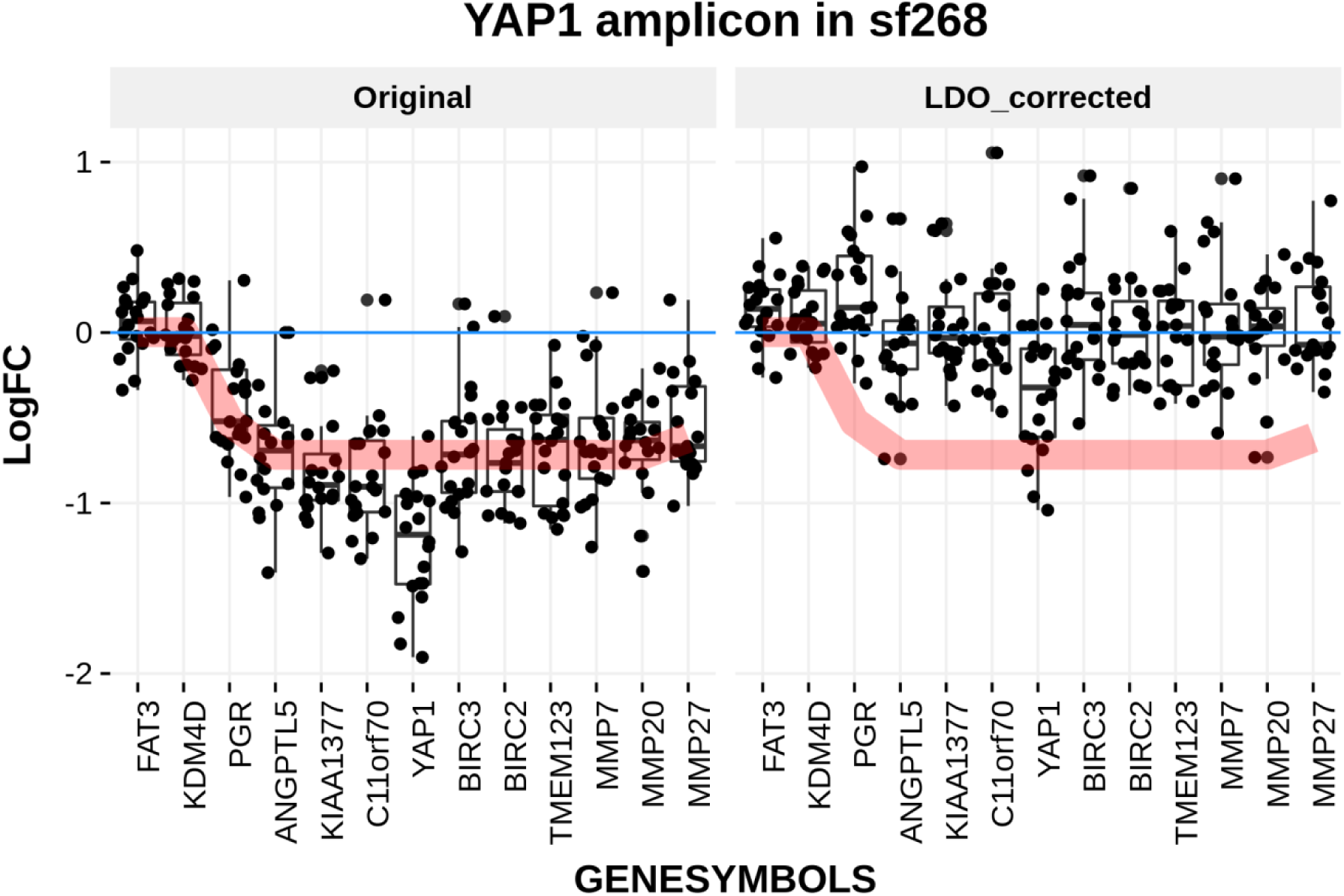
YAP1 Specific LDO correction. Sensitivity, calculated as LogFC, conferred by each guide (black dots) within the YAP1 amplicon in the sf268 cell line summarized by a boxplot for each gene in the amplicon. To ease the interpretation, the red line displays the inverted copy number value scaled to the data. The left panel displays the uncorrected sensitivity scores, while the right panel shows the sensitivity scores after LDO correction.

In Fig. 2 b), the resulting corrected sensitivity scores are shown in the *YAP1* amplicon in sf268. The copy number effect has been successfully removed and *YAP1* still scores as significantly lethal, thereby being identified as the amplicon’s driver, as is expected from existing shRNA screens and reported elsewhere (11,12).

Overall, LDO removes the copy number effect beyond the *YAP1* amplicon in sf268 and in mkn45 cell lines, as shown in Fig. 3. The number of guides with log2(CNA) larger than 2 and LogFC below -0.5 is decreased from 84 to 25 guides in mkn45 and 274 to 50 guides in sf268.

**Figure 3.**
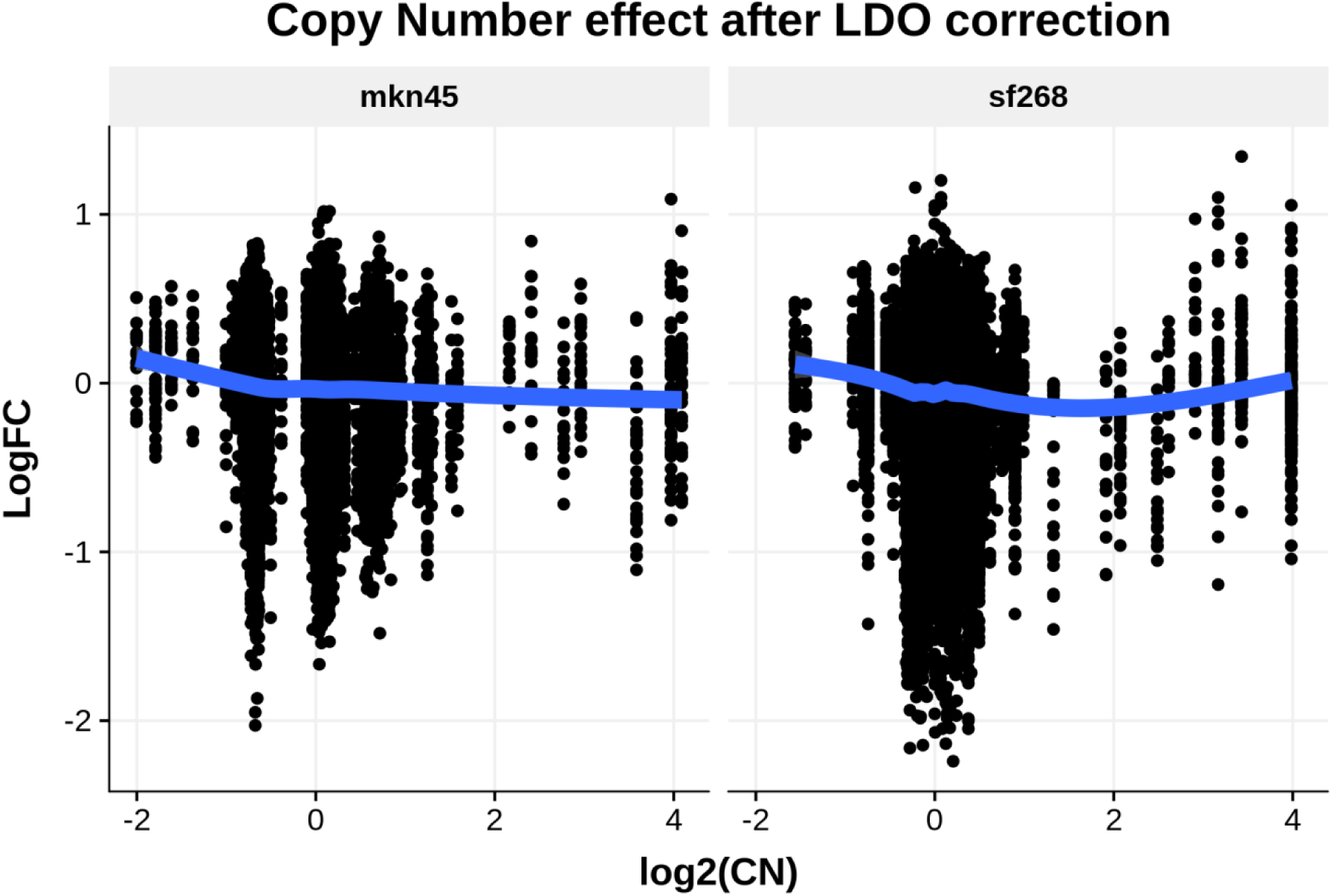
Global LDO Correction. The sensitivity to CRISPR-mediated knock-out after LDO correction is not dependent on the level of DNA amplification of the underlying genomic region anymore.

### Library design and guide quality

Although the method applied on this screen was able to successfully recover the driver in the *YAP1* amplicon, this is not always the case as shown in Fig 4.

**Figure 4.**
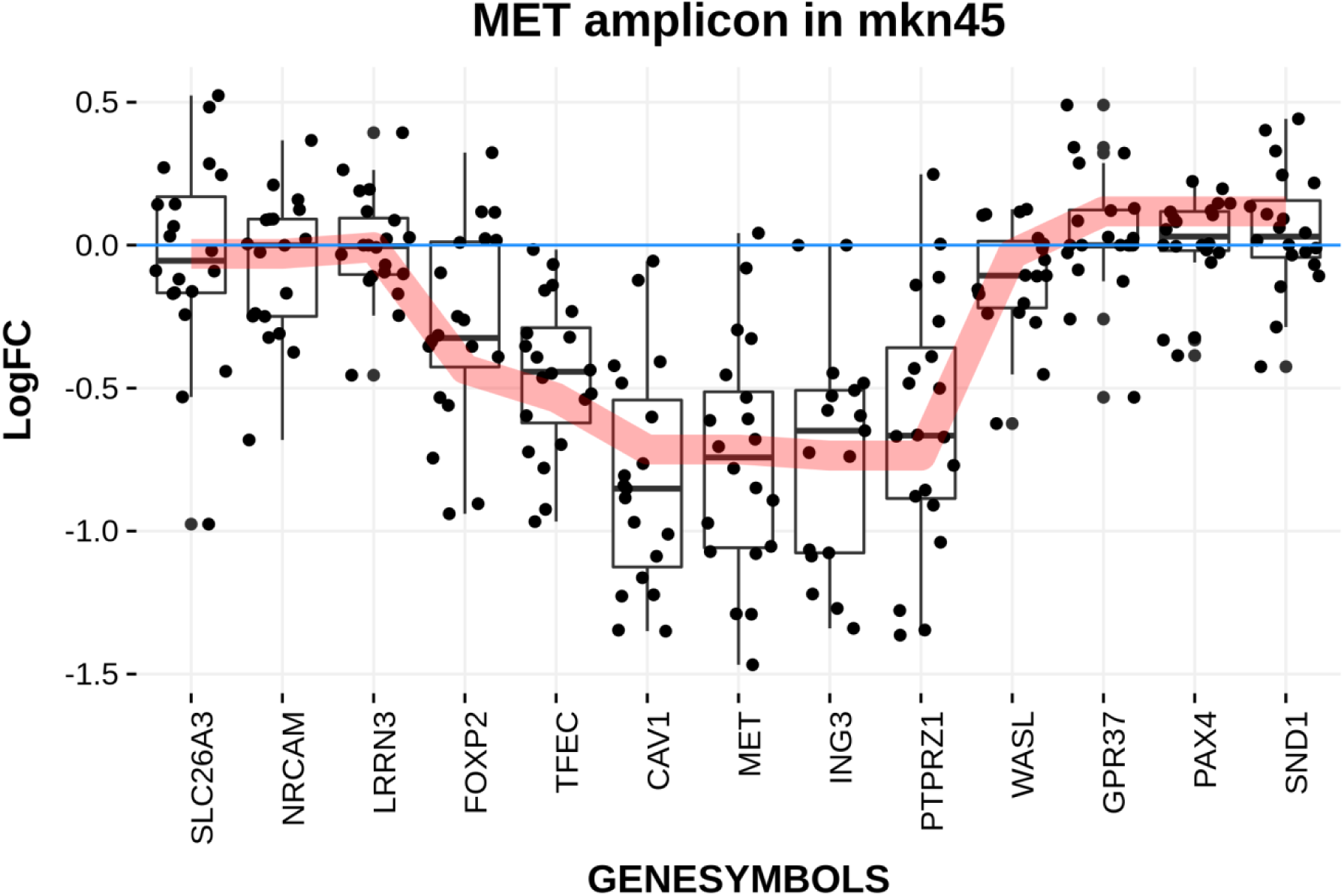
MET Specific LDO correction in mkn45. Sensitivity conferred by each guide (black dots) within the MET amplicon in mkn45 summarized by a boxplot for each gene in the amplicon. The red line displays the inverted copy number value scaled to the data.

From shRNA screens and other reports (13,14), *MET* is expected to be the driver of this amplicon. Therefore, one could expect the *MET* guides to display a stronger relative drop out compared to the rest of the genes in the amplicon. However this was not the case and thus applying the LDO correction did not enable the recovery of *MET* as the driver of the amplicon. The degree of amplification does not appear to explain the lack of differential *MET* effect in mkn45 considering that the amplification in sf268:*YAP1* is equivalent to what is seen in mkn45:*MET*.

One potential reason for this lack of relative drop out is the quality of the guides used. The screen was rerun with different guide designs. The result for the *MET* amplicon in sample mkn45 is shown in Fig. 5 and in this case, *MET* does display a stronger phenotype than the rest of the amplicon. This highlights the need for careful library design (Supp. Fig. 1).

**Figure 5.**
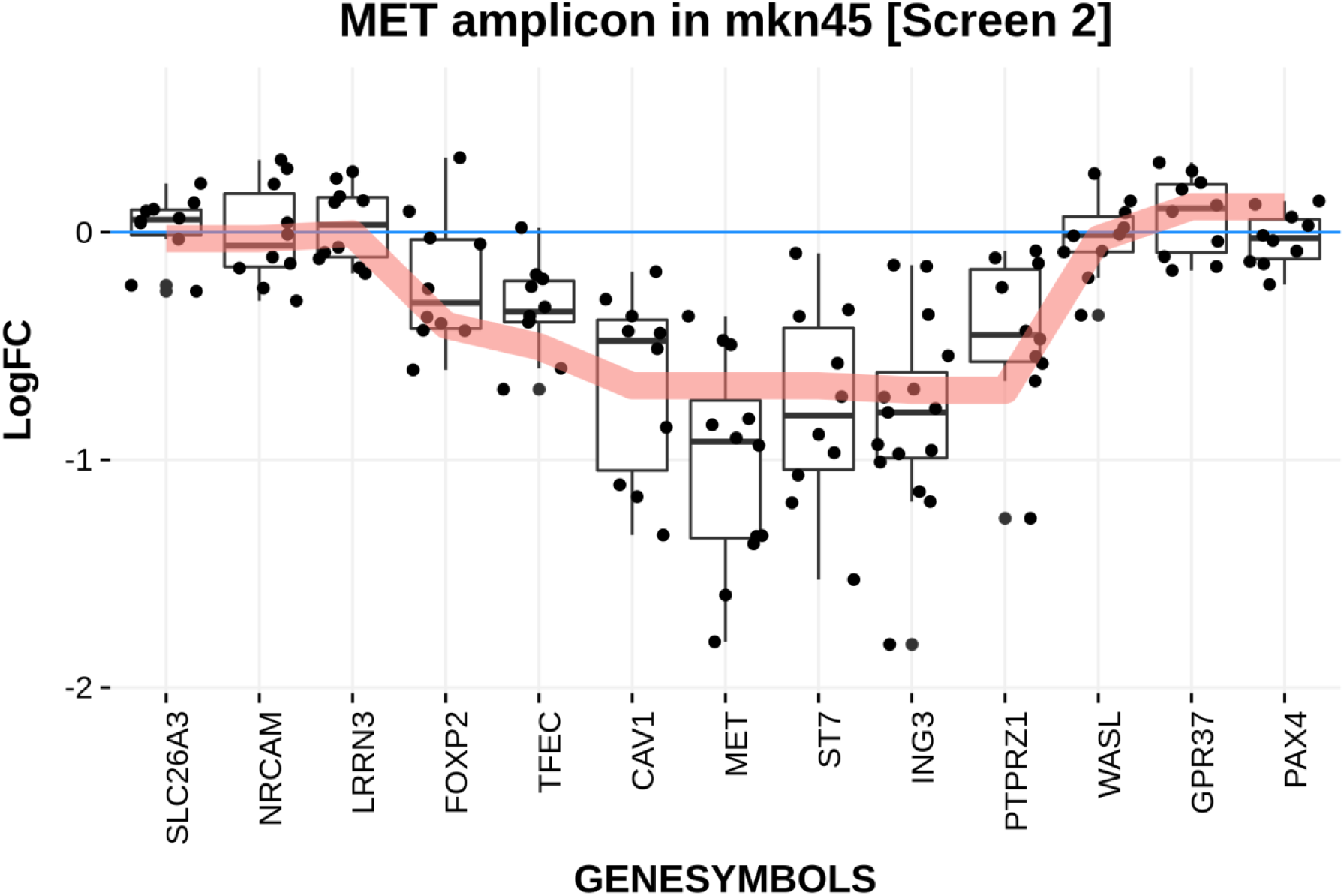
MET Specific LDO correction in mkn45’s second screen. Sensitivity conferred by each guide (black dots) within the MET amplicon in mkn45 summarized by a boxplot for each gene in the amplicon. The red line displays the inverted copy number value scaled to the data.

### Generalized Additive Model (GAM) method

The LDO method proposed above does not rely on any orthogonal data, such as copy number, and can even be used to estimate the copy number of the screened samples (see Supp Fig 1). However when available, the measured DNA copy number values can be used to adjust the sensitivity scores. To do this, we used a generalized additive model (GAM, (15)) framework and modelled the sensitivity to CRISPR-mediated gene knock-out as a function of copy number to yield an adjusted CRISPR-mediated gene knock-out estimate. The potential benefit of this framework compared to LDO is that it can be extended to consider any arbitrary number of additional features (both linked to artefactual or true effects) potentially relevant for the purpose of modelling the phenotype (e.g. gene expression, multi-alignment of guides, etc.). In this analysis, only the copy number measurements were used. Once the model has been fitted, the effect of the artefactual components of the sensitivity can be removed from the observed phenotype in order to keep the “biologically-relevant” component (in this example only the artefactual copy-number effect is considered and removed). Unlike the LDO method, the GAM method is insensitive to the screen density and would be preferred should a sparse coverage of the genome be considered in the screen. Additionally the GAM method does not require a prior list of known essential genes to be performed.

The GAM structure can be written as follows:

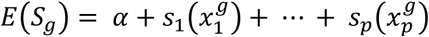

Where *E*(*S_g_*) is the expected sensitivity of guide 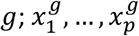 are the predictor variables for *g* and 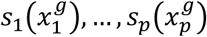 denote the smoothing functions estimated by non-parametric means from the data. Finally, *α* is the intercept. Note the lack of a linker function in the above equation compared to the canonical GAM framework, since we consistently use the identity function. For the purpose of fitting the GAM to the data, we use the R implementation from the mgcv package (16) with default parameters, so that penalized thin plate regression spline models are used for the smoothing.

This framework enables us to take into account an arbitrary number of predictor variables to model both linear and non-linear dependencies of the data. The aim is to remove from the measured sensitivity *S_g_* the components of the model, which are deemed to come from artefactual predictor variables (e.g. copy number) but keep those coming from variables which are considered true predictors of biological sensitivity (e.g. gene expression). Instead of 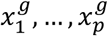 let us further differentiate the predictor variables into the artefactual variables 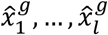 and the explanatory variables 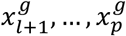 so that the GAM corrected sensitivity can be written as

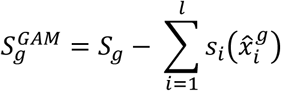

In this presentation a single artefactual predictor 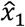 is used which represents the copy number value at the position of guide *g*. The GAM corrected sensitivity score 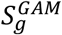 can then be used in lieu of the original sensitivity score with the same hit-defining thresholds and interpretation.

The correction of the copy number artefact in sf268 and mkn45 using GAM is shown in Supp. Fig. 2.

Additionally the corrections based on LDO and GAM in the genomic regions with measured copy number values are consistent as shown in Supp. Fig. 3.

## Discussion

The use of high-throughput CRISPR screens to identify cell autonomous cancer dependencies has become routine. However, as shown in previous studies, these screens display high rates of false positive hits in regions of high copy number amplifications. In this report, we describe two methods, Local Drop Out (LDO) and General Additive Model (GAM), to correct for this copy number bias, thereby enabling the identification of true positive hits while reducing false positives substantially. In both cases the methods were developed with experimental setups in mind utilizing only a few number of cell lines, including single model experiments. Thus making these methods appropriate for a broad range of experiments. As a result the CN artefact corrections proposed are performed at the level of single samples. We applied both methods to previously published screening data of 2722 genes performed in the sf268 and mkn45 cell lines. The utility of the methods were shown by way of two examples: first, the *YAP1* dependency in sf268 was recovered, while removing 8 false positive genes from the hit list (*ANGPTL5*, *KIAA1377*, *C11orf70*, *BIRC3*, *BIRC2*, *TMEM123*, *MMP7*, and *MMP20*); second, the *MET* dependency in mkn45 was recovered in one of the two screens, while removing 3 false positive hits (*CAV1*, *ST7*, and *ING3*). Overall, the number of guides with log2(CNA) larger than 2 and LogFC below -0.5 is decreased from 98 to 29 guides in mkn45 and 267 to 41 guides in sf268 when using LDO; with GAM the number of guides are reduced to 28 and 37 guides for mkn45 and sf268 respectively.

These methods, however, do have limitations. We observed that rescuing true positives within amplicons is only possible if the driver mutation in the amplicon of interest is indeed displaying a stronger relative drop out than the neighbouring genes. Depending on the guides used, this is not always the case as demonstrated with *MET* in mkn45 in our first screen. Despite this caveat both methods are still able to remove false-positives, although the true positive is not rescued. We would argue that in a typical screening effort, the loss of a few true positives is less damaging than the large amount of false positives proposed by the first alternative of using the uncorrected scores. Indeed, a lot of effort and resources can be spent chasing an elusive false positive. The second obvious alternative is to remove any amplified region from the subsequent analysis, which means a large amount of false negative hits, since those would not even be considered for further analysis, but also relies on prior available copy number measurements which is not always the case.

Another limitation is that these methods are highly dependent on guide scores obtained in the screen which can be variable. In our mkn45: *MET* example, it is unclear what the reason for the difference in drop out of the driver is. Guide design could be an explanation, however if CRISPR genome editing does indeed generate two cellular responses in cancer cells as suggested in (6): an early anti-proliferative DNA damage response and a later gene dependant effect, the number of doublings before harvesting could also be an explanation. Whereby cell lines with long doubling times would only undergo enough doublings to sustain the DNA damage response but not enough to signal a differential effect from the driver genes. This hypothesis however does not seem to fit with the doubling times of 29h and 44h for mkn45 and sf268 respectively as reported in the Cancer Cell Line Encyclopedia (CCLE) (17).

Outside of these limitations, each of the two methods presented offer different advantages to the correction for the copy number induced false positives in loss-of-function CRISPR screens. The LDO method can correct for the copy number artefact even when copy number is not known beforehand as long as the density of the CRISPR screen is high enough to capture the copy number events with confidence. On the other hand, the GAM correction method requires copy number measurements, but it is not dependent on high density screens and can additionally incorporate an arbitrary number of predictor variables in its model. The fact that LDO does not need any copy number information also enables the user to estimate the underlying copy number of its sample by exploring the magnitude of the correction that was applied to the different genomic regions (Supp. Fig. 3).

## Material and Methods

### Essential Genes

To collect an initial list of essential genes the results from (18) was used. In particular the essentiality of each gene was established in (18) using a genome-wide single guide CRISPR screen in 4 cancer cell lines. The strength of the essentiality is reported as an adjusted p-value in the accompanying data. Here, the genes with a maximum adjusted p-value of 0.05 across all 4 cell lines are used as de facto essential genes if and only if the accompanying CRISPR score is also smaller than -1. This results in a list of 814 potential essential genes (Supp. Table 1).

### LDO

The choice of the exponential decay function in the weighted mean calculation is arbitrary (and any weighing function can easily be used instead in the provided scripts). Any monotonously decreasing function could be used, or, for example, a simple sliding window. The size of the window, or the value picked for *ω* in the exponential decay case, should be chosen so as to borrow the information from as many genes as possible while still remaining within the bounds of the expected copy number event sizes that are expected to be observed. The exponential decay function has the advantage of putting more weight to the genes in the direct neighbourhood of the gene of interest and thus even if the size of the window considered is relatively large the estimate remains relatively robust.

Similarly the values for *α*_1_, *β*_1_, the minimum number of three genes per short CN event and the choice of only considering events with correction values larger than the 1.5 times the median average deviation of the background noise were chosen arbitrarily based on a priori expectation of the effects we wish to correct for.

